# Implementation of an Open Source Software solution for Laboratory Information Management and automated RNAseq data analysis in a large-scale Cancer Genomics initiative using BASE with extension package Reggie

**DOI:** 10.1101/038976

**Authors:** Jari Häkkinen, Nicklas Nordborg, Olle Månsson, Johan Vallon-Christersson

**Affiliations:** Lund University, Department of Clinical Sciences Lund, Division of Oncology and Pathology, Lund, Sweden.

**Keywords:** Next-generation sequencing, RNAseq, Genomics, Software solution, BASE, LIMS, Laboratory Information Management System, Analysis pipeline, Breast cancer

## Abstract

**Background:** Large-scale cancer genomics initiatives and next-generation sequencing for transcriptome profiling allow for detailed molecular characterization of tumors, and provide opportunities for clinical tools to improve diagnosis, prognosis, and treatment decisions. Laboratory information, data management, and data sharing in large-scale genomics projects is a challenge. Aiming to introduce such technologies in a clinical setting offer additional challenges associated with requirements of short lead-times and specialized tracking of biomaterials, data, and analysis results.

**Results:** Using the free open-source BioArray Software Environment (BASE) and extension package Reggie we have implemented a laboratory information management system and an automated RNAseq data analysis pipeline that successfully manage a large regional cancer genomics initiative. The system manages enrolled cancer patients, tumor biopsies, extraction of nucleic acid, and whole transcriptome RNA-sequencing through to data analysis and quality control. The implementation offers integration of laboratory equipment and operating procedures, and information tracking in a module based fashion enabling efficient and flexible use of personnel resources. The system provides two-factor authentication and transaction control and seamless integration of freely available software for RNAseq analysis such as Tophat, Cufflinks, and Picard. As of February 2016 more than 8000 patients and over 6000 tumor biopsies have been successfully processed. Lead-time from biopsy arrival to summarized reports based on RNAseq data is less than 5 days, in line with regional clinical requirements. BASE and Reggie are freely available and released as open-source under the GNU General Public License and GNU Affero General Public License, respectively.

**Conclusion:** Using free open-source software together with BASE and a customized extension package, Reggie, we have implemented a system capable of managing large collections of quality controlled and curated material for use in research and development and tailored to meet requirements for clinical use. Featuring high degree of automation and interactivity the system allows for resource efficient laboratory procedures and short lead-times with demonstrated use of RNAseq data analyses in a clinical setting.

## Background

When microarrays were first introduced in the mid 1990s, users were initially faced with challenges of managing and analyzing the unprecedented large amount of data generated from even relatively few experiments. Over the past decade, next-generation sequencing has emerged with increased amount of data output that again challenges bioinformatics and data management. Gradually, since their introduction, these technologies have matured and become more readily available. In parallel, numerous tools and systems were developed to meet the accompanied challenges of data management and analysis [1]. A large part was, and still is, developed within the research community as open source software projects that allow for collective efforts and integration of available software within new systems [2–4]. Increasing availability and decreasing cost, in combination with the possibilities offered, have lead to widespread use of these technologies in many research projects. Early omics-projects included detailed molecular characterization of tumors with the aim of identifying new cancer biomarkers and introducing clinical tests based on microarray or sequencing assays [5, 6]. Successful development and validation efforts require large sample sets meticulously controlled for selection and technical biases. Furthermore, for transition to clinical use to be practically and economically viable, laboratory processes, information management and analysis need to be time and resource efficient to meet requirements dictated by clinical practice. To this end, management and analysis of data generated by omics-technologies remain a challenge. To meet these challenges we implemented a laboratory information management system and an automated RNAseq data analysis pipeline for a large regional cancer genomics initiative in the southern health care region of Sweden [7]. During the implementation process, the goal was to use freely available tools and to develop software that are made freely available and released as open-source under the GNU General Public License family of licenses.

## Implementation

Our implemented system uses a standard BASE [8, 9] installation and Reggie; a customized extension package that serves as a streamlined interface with tools designed for registration of laboratory information, generation of protocols, as well as execution of analysis and retrieving reports based on RNAseq data (Figure 1).

The system integrates external file servers for storage of data that is not imported into the BASE database and computation clusters based on Rocks Clusters [10].

**Figure 1.**
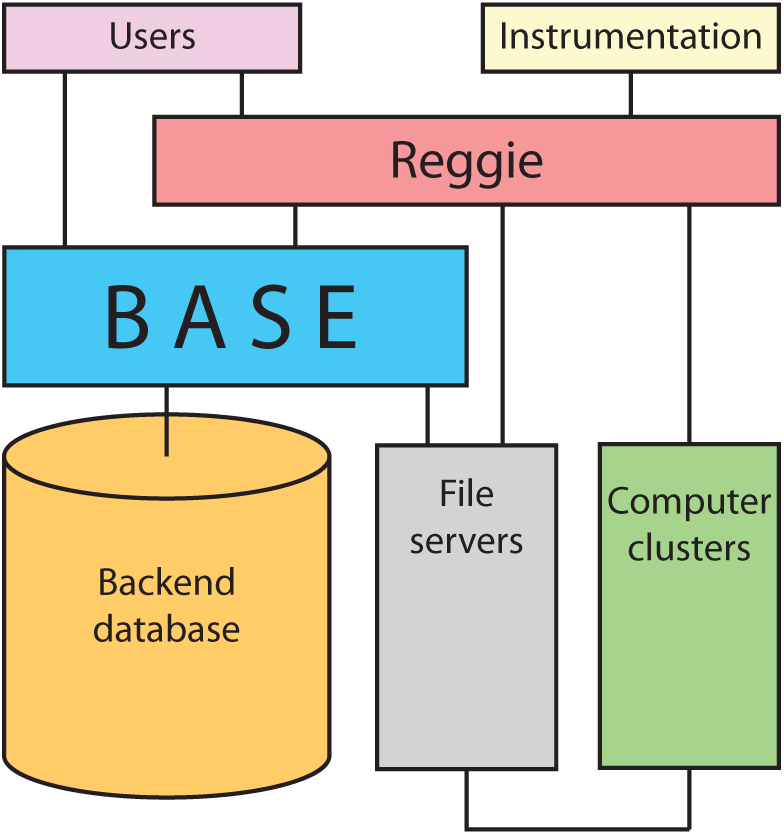
Schematic overview of the implemented solution for laboratory information management and automated RNAseq data analysis. At the center of the implementation is a standard BASE installation (blue) with a back-end relational database (orange). Extension package Reggie (red) is designed as a core controller and logic layer that holds information about laboratory operating procedures. Users (purple) with individualized permissions can interact directly with BASE or through Reggie wizards that provide interfaces for registration of information and integrate laboratory instrumentation (yellow). Importantly, Reggie wizards can execute complex workflows and the wizards control, resolve and establish logical relations in the database through BASE. Reggie can run external tools and queue jobs on computation clusters (green) and Reggie monitors and verifies multi-step analysis pipelines. For this purpose, Reggie can access, retrieve and modify information directly on the file servers (grey) or through BASE. Interactions performed directly by Reggie are still registered in BASE.

The basis of the implemented system is a minimalistic but complete model designed as a representation of the major parts of the laboratory process; patients, cancer diagnosis, biopsies, biomaterial lysates, necessary extracts, library preparation, library pools, sequencing, and required derived data items (Figure 2a).

**Figure 2.**
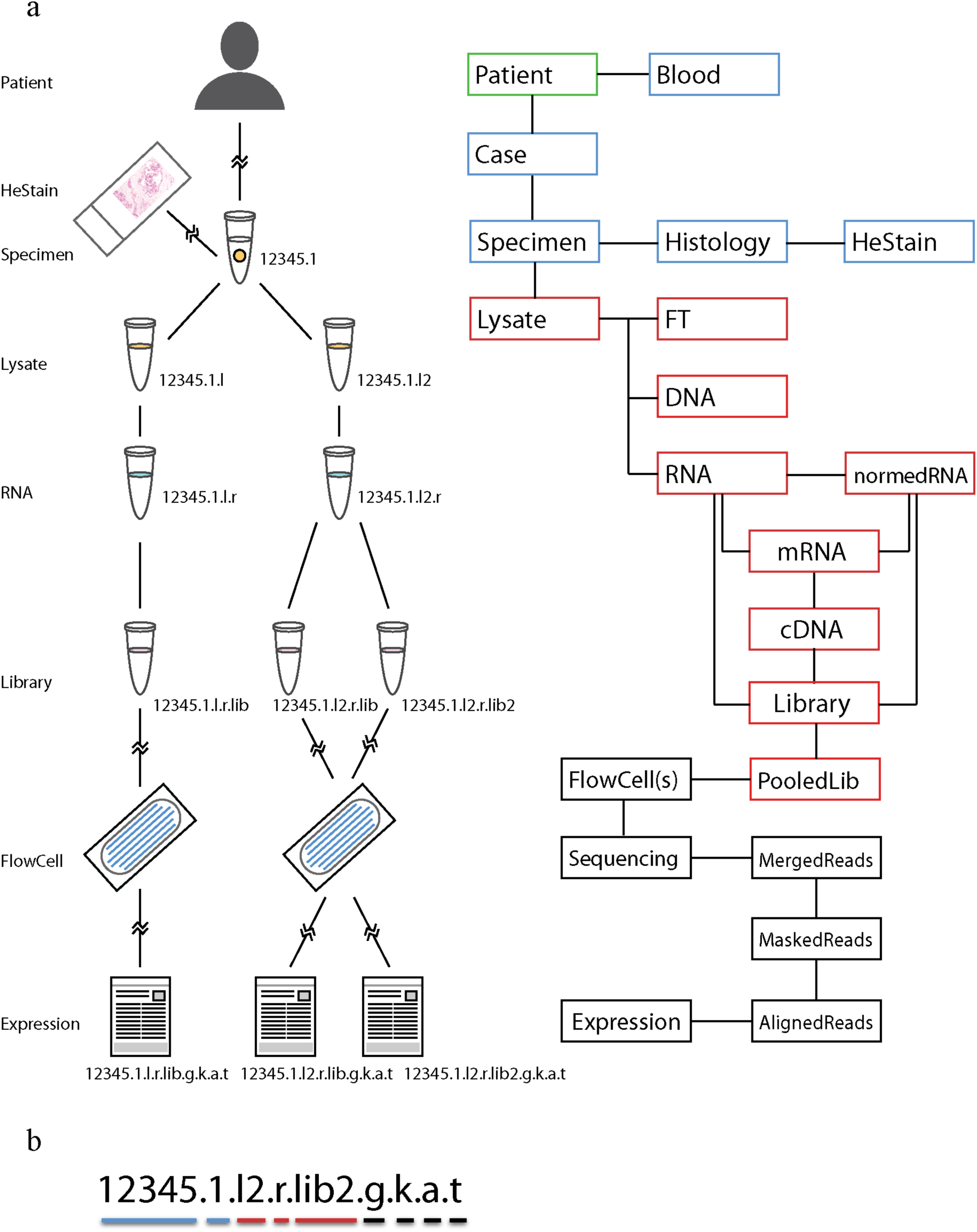
Laboratory process and corresponding model with naming structure for included database items. (**a**) Genealogy for a tumor RNAseq experiment (left) and the implemented model with database items (right). The cartoon flowchart (left) illustrates an example of a patient and laboratory procedure for a received tumor biopsy with names for database items in accordance with the used naming structure. Relational connections in the cartoon are illustrated with lines or broken lines for intermediate steps that are omitted from the illustration. The implemented model (right) is illustrated as a flowchart with colored boxes. The model is designed as a representation of the major parts of the laboratory process; patients (Patient), cancer diagnosis (Case), tumor biopsies (Specimen), sample lysates (Lysate), extracts including RNA (RNA), sequence ready libraries (Library), library pools (PooledLib), flowcells for sequencing (FlowCell), sequencing (Sequencing), required derived data items such as demultiplexed and merged reads (MergedReads), and derived gene expression estimates from RNAseq data (Expression). The complete model includes a number of additional database items (not shown), as well as representations for blood samples (Blood), tumor biopsy part used for FFPE (Histology), and concentration normalized RNA aliquots (normedRNA). (**b**) The logical naming structure is constructed using strings separated by periods and read from left to right. The example shown is a name for the database item representing derived gene expression estimates (Expression). The first two name parts (blue underline), make up a unique specimen tube ID, i.e., 12345.1. Following the specimen tube ID, additional suffixes are appended to represent sequentially derived items from the laboratory and analysis process, e.g., suffix “.l” for lysate, suffix “.r”, for extracted RNA, and suffix “.lib” for prepared sequence library. A split in the lineage, for example partitioning of a biopsy into two separate parts that are individually lysed in preparation for nucleic acid extraction, is handled by adding a serial number to the derived item suffix, that is “.l” and “.l2” for the first and second lysate respectively (see example in Figure 2a and 2b). Three name parts represent the lineage through laboratory processing items (red underline): lysate (.l2), RNA (.r), and library (.lib2). The last four name parts (black underline) capture secondary analysis items: demultiplexed and merged reads (.g), masked reads (.k), aligned reads (.a), and derived gene expression estimates (.t).

In designing the model, careful considerations were made to include database items for physical entities that need to be tracked in the laboratory, such as extracted RNA and other biomaterial aliquots, while avoiding redundancy by excluding items for many intermediate biomaterials and laboratory steps. However, included items are used as placeholders for associated properties, protocols, and events, such as creation events, and this allows for tacking of procedures and biomaterials omitted from the model. A deliberate effort was also put into conceiving a logical naming structure for included database items that allows for names to hold information about relations between items in human understandable format (Figure 2a and b). This was done to allow relational information about individual items to be directly available when data is exported from the database, e.g., when item names are listed in spreadsheets or printed on tube labels. The naming scheme also helps in understanding the data structure and relations when directly interacting with data stored on external file servers. Whereas the model and relational database store information and maintain logical connections between items, the extension package Reggie is designed to provide data exchange interfaces to assist in controlled information input and output to ensure database integrity. As such, Reggie is the core controller and logic layer that holds information about laboratory operating procedures and can execute complex workflows in a context sensitive manner. By utilizing Reggie wizards (Figure 3), laboratory procedures can be modularized and many tasks can be performed asynchronously without undesirable consequences of race conditions and database inconsistencies. The Reggie extension reduces lead-time, errors, and simplifies user interaction with instrumentation during the laboratory work by integrating laboratory equipment. The level of integration depends on hardware capabilities where in some cases direct transfer of data into the laboratory information management system (LIMS) is achieved, whereas in many cases exchange of configuration files to control laboratory equipment and result files for parsing and information retrieval are required. Reggie wizards are also designed to execute and monitor multistep analysis pipelines that include automatic initiation of demultiplexing of data from pooled sequencing runs, quality control, alignment, and, for RNAseq, summarization of gene expression values.

**Figure 3.**
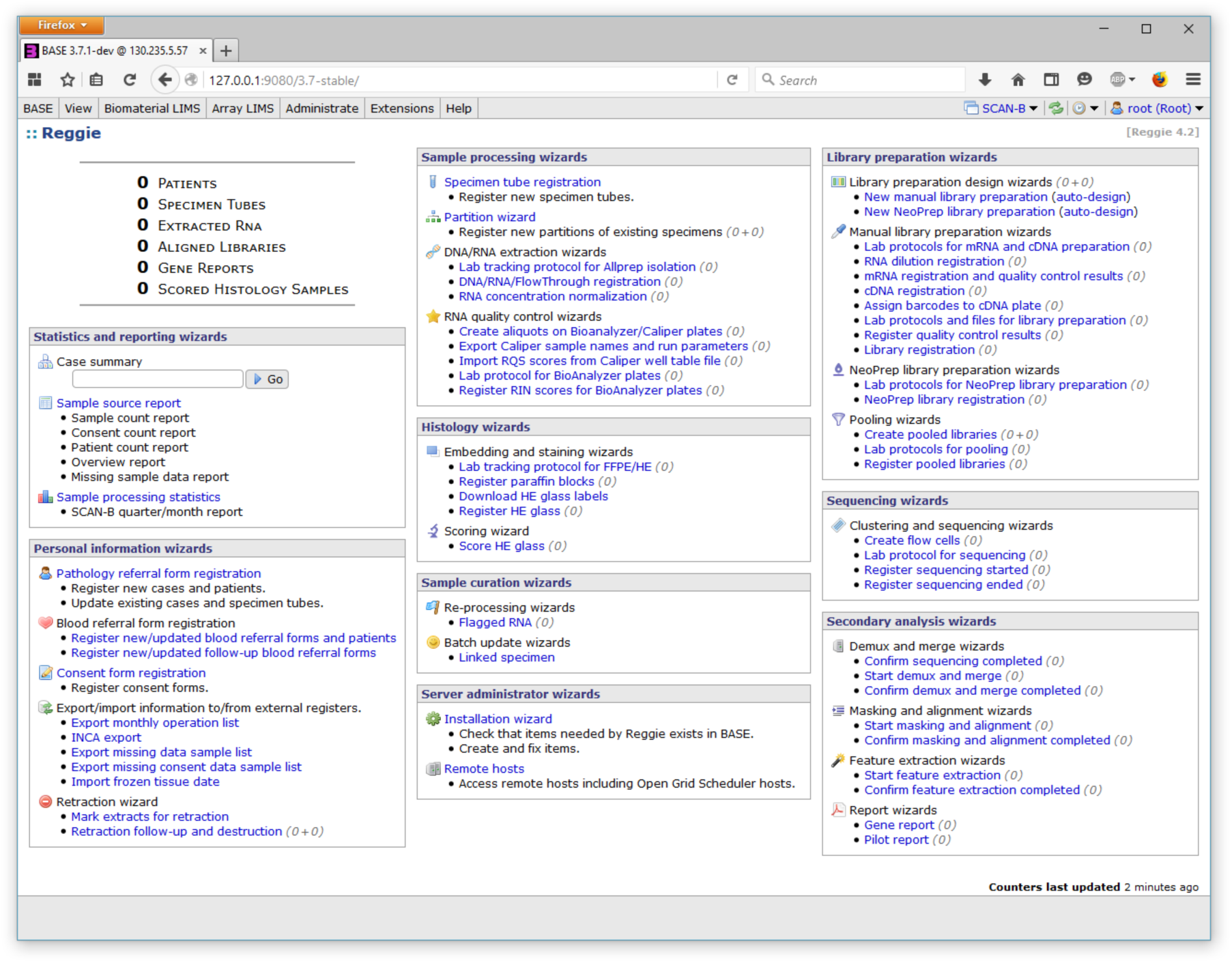
Main page of Reggie extension package for BASE. The main page of extension package Reggie provides access to implemented tools, or wizards, to support tasks in the laboratory and data analysis pipeline. Customized wizards are also included for generating database statistics or managing patient information and more. In addition, the main page includes counters (top left) with totals for number of items registered at an installation instance of the BASE/Reggie set up. Numbers within parenthesis after links to Reggie wizards are counters for items pending further processing by respective tool. The example screen shot is from a pristine installation and all counters are zero.

## Results and discussion

### BASE 3

To efficiently manage sequencing based experiments, major updates were made in the 3^rd^ major release of BASE. The first generation BASE [8], released in 2002, was designed for microarray laboratories as a data management and analysis system. It featured a multi-user local data repository with a web browser based user interface and included a laboratory information management system for biomaterials and array production, annotations, hierarchical overview of analysis with capability to integrate external analysis tools. Continued development has introduced major changes, such as a reimplementation written in Java and using Hibernate for object/relational persistence in the back-end relational database management system and support of storage of data in files rather than solely relying in storing all data in database tables depending on data [9]. However, for historical reasons the database schema remained tightly linked to procedures in microarray experimentation, which imposed limitations for flexible use in sequencing projects. Therefore, the current and third generation BASE introduced more generic database items to support complex models for sequencing experiments and an updated extensions system with interface points provided by the BASE core API to permit development of more elaborate extension packages. Example extension packages are Reggie, described herein, and LabEnv, an extension that integrates networked temperature and humidity recorders placed in the laboratory for continuous and automated collection of environment data. Another example is the YubiKey extension that adds support for two-factor authentication using personal YubiKey one-time-password sticks. All extensions are available from the BASE plug-ins site [11]. Data and items stored in BASE can be organized in projects and shared between groups and users with great detail. Items can be listed in customizable table views along with properties and associated information. Table views can be filtered, sorted and used to export included information as tab-separated text or XML files. In addition, dynamic lists have been introduced that permit filters to be applied on database items at multiple levels in the genealogy. Furthermore, BASE supports configurable storage plates to organize and track physical storage location, as well as reaction plate items to represent laboratory steps performed in microtiter plates. Detailed description of features and development is available at the BASE project page [12].

### Reggie

Reggie is an extension package designed as a view and control layer that takes full advantage of the core API enhancements made available in BASE 3. The extension is tailored to the laboratory process and laboratory operating procedures used in a regional breast cancer genomics initiative in the southern health care region of Sweden and includes several tools, or wizards, to support tasks in the laboratory and data analysis pipeline (Figure 3).

Wizards are also included for generating statistics and managing patient information such as pathology referral form registration. In addition, Reggie wizards guide users through the required steps of removing biomaterial and information in the event that an enrolled patient retracts informed consent. Importantly, Reggie is designed to handle a number of operations independently that otherwise typically need to be done in a certain order to avoid undesirable consequences of race conditions. For example, Reggie can manage asynchronous registration of items with logical connections in the database such as a cancer patient, tumor diagnosis and associated tumor biopsies (specimens). This is especially useful in large projects where it is desirable to modularize tasks so they can be executed in parallel. Specifically, operating procedures are designed to accept registration of personal patient information by a patient curator. Meanwhile, laboratory personnel can register tumor biopsies independently as soon as they arrive in order to start sample processing without delay. During these operations, Reggie controls, resolves and establishes relations between asynchronously created items in the database. Conversely, successive laboratory steps need to be performed in sequential order, and Reggie wizards are designed to facilitate modularization of workflows so that one person can perform one step and seamlessly hand over the subsequent step to another person. Similarly, Reggie can start external tools and queue jobs, monitor progress and auto-confirm and execute subsequent steps when preceding ones are successfully completed. In this manner, multi-step analysis pipelines are automatically executed as well as automatically triggered by the completion of a sequencing run. Thus, significantly reducing manual intervention and overall lead-times. While the extension package Reggie includes tools tailored to the needs in a specific cancer genomics initiative, many are nonetheless designed for standard laboratory operations. In the following sections, a number of Reggie wizards in the RNAseq pipeline are described in more detail to demonstrate implemented solutions for workflows commonly used in genomics projects in general and to highlight the possibilities when using a customized extension package with BASE. A comprehensive description of Reggie features and development is available on the BASE plug-ins page [11].

#### Personal information wizards

The implemented system takes advantage of the multi-user functionality of BASE where individual users can be assigned roles or group memberships configured with different permissions. In this manner, different users can be given personalized access to Reggie wizards. In addition, patient items stored in the system have generic and pseudonymous patient identifiers where sensitive information is masked for most users. For example, only users that are patient curators have access to personal information wizards and private information stored in the database. The personal information wizards comprise a set of customized tools for entering sensitive information received from referral forms and clinical professionals, and for exporting personal data used to request follow-up data from regional cancer registers. It also includes a set of wizards for registering informed consent from patients or for executing work-flows that mark, and subsequently remove, database items in the event that an enrolled patient choose to revoke a consent.

#### Sample processing wizards

Sample processing wizards include tools for specimen tube registration through partitioning of biopsies and extraction of nucleic acids and RNA quality control. An integral part of operating procedures in the laboratory is a unique barcoded identifier on the referral form that accompanies received specimen tubes. The sample processing starts by registration of specimen tubes and scanning this identifier. Multiple tubes can be registered from the same referral form and Reggie will assign unique names to each tube.

#### Specimen tube registration

Reggie provides dialogs for entering referral form identifier and associated information received on the referral form, such as sampling date and time, laterality for breast cancer diagnosis, and biopsy type, e.g., biopsy from surgery specimen or needle core-biopsy. During the registration, Reggie will assign storage location for received tubes by retrieving suitable free physical locations, i.e., storage box, box coordinates, and freezer location. Once registered, specimen tubes will appear as a list of tubes in the partitioning wizard pending further processing.

#### Partitioning wizard

The partitioning wizard provides a user interface for partitioning of received biopsies into separate parts: one for nucleic acid extraction, one for preparation of a formalin-fixed and paraffin embedded (FFPE) portion, as well as a remaining part to be stored if any remains. The wizard executes a workflow that creates necessary database items and to facilitate laboratory work, creates a label file for easy printing of barcoded tube labels. The interface integrates a laboratory scale to directly retrieve weight of individual biopsy portions and save these as properties of respective database item. After completion of the partitioning process, lysate items representing the biopsy portion that will undergo lysis and homogenization in preparation of nucleic acid extraction, show up in the DNA/RNA extraction wizard pending continued processing, whereas histology items intended for FFPE show up in the histology wizard.

#### DNA/RNA extraction wizards

Reggie provides procedures for DNA/RNA extraction using QIAGEN Allprep spin-column kits with automated sample prep on the QIAcube (QIAGEN). The Reggie workflow for extraction is designed to permit selection of 1 to 12 lysates pending processing and will configure a QIAcube run accordingly. Furthermore, a downloadable tracking sheet intended for use in the laboratory is created that includes assigned positions to correctly load the QIAcube centrifuge. When executed, the Reggie wizard will create items for extracted RNA/DNA and flow-through aliquots and assign physical storage locations, as well as aliquots that will be used for RNA integrity control on a PerkinElmer Caliper LabChip station (PerkinElmer). The wizard also presents an interface for integrated concentration and purity measurements of extracted nucleic acid using NanoDrop 8000 UV-Vis Spectrophotometer (Thermo Scientific). RNAs that are extracted with sufficient yield for sequencing, according to criteria in the operating procedures, are automatically flagged for concentration normalization. In doing so, Reggie will present the user with instructions of how to prepare normalized RNA aliquots for direct input in downstream protocols for preparation of sequencing libraries. To expedite the workflow while minimizing errors, the instructions come complete with amount of RNA to use based on concentrations stored in the database.

#### RNA quality control wizards

A collection of wizards is provided to assist in configuration, running, and registration of results from assays assessing RNA integrity. Quality control aliquots from the DNA/RNA extraction are automatically assigned wells in microtiter plates for analysis with Caliper in accordance with operating procedures. Configuration files for running Caliper are created by Reggie for upload to the Caliper controller software. Subsequently, Reggie receives results files that are parsed and analysis results are imported into BASE where data is associated with respective RNA aliquot items. A wizard for manual configuration of Bioanalyzer runs is also available to facilitate additional quality control runs if required. Information from RNA quantification and quality control is stored as properties and annotations to database items and are available in the downstream workflows for library preparation and sequencing.

#### Histology wizards

A highly specialized set of wizards is included for managing FFPE preparation, sectioning, and hematoxylin and eosin (HE) staining, as well as immunohistochemistry. A scoring wizard is included to provide an interface for histological review of stained sections, facilitating input of a set of predefined cellularity scores that are useful in evaluation of cancer genomics results. The scoring wizard also includes a user-friendly drag-and-drop interface for uploading digital images captured by the microscope software. Images are associated with the respective histology item and readily available when reviewing data or for creating reports.

#### Library preparation wizards

Library preparation wizards include tools for laboratory steps in preparing sequencing ready pooled libraries from RNA through mRNA purification, assignment of barcodes, quality controls, and pooling libraries for multiplexed sequencing runs. As with sample processing, operating procedures dictated the steps implemented in the Reggie workflow.

#### Library preparation design wizards

Sequencing libraries are prepared in batches that are subsequently pooled for sequencing in multiplexed runs. Reggie tools provide an elaborate interface for selecting available RNA aliquots or pre-normalized RNA aliquots and configure these in batches on microtiter plates for manual library preparation or with a NeoPrep Library Prep System (Illumina) for automated library preparation. Presets for plate layouts are available in accordance with laboratory operating procedures and the selected plate configurations can be populated with available RNA aliquots manually or by auto-select functionality where RNAs pending library prep are automatically chosen according to criteria defined in the Reggie workflow. Once a library preparation design is registered in Reggie, a context specific lab-tracking sheet is created and available for laboratory personnel to proceed with following steps in the procedure. Instructions in the created lab-tracking sheet depend on the library preparation design, for example, if dilution is needed, calculated recipes based on specified amounts and stored concentrations are provided together with storage locations of included RNA aliquots. If, on the other hand, the library preparation design comprises pre-normalized aliquots, dilution recipes are not needed nor included. Importantly, designs using combinations of both examples are supported, and the lab-tracking sheets are devised with the overall aim of reducing time consumption and risk of errors when working with library preparation.

#### Assign barcodes to cDNA plate and Lab protocols for library preparation

A separate wizard is designed to assign barcodes to libraries. Depending on the previously defined library preparation design, which may include one or more sets of libraries intended for pooling, presets of compatible selections of barcodes are available. Manual selection of barcodes is permitted and the user will be notified if incompatible combinations are selected. Analogous to lab-tracking sheets for setting up library plates, context sensitive protocols are created to assist in the laboratory steps of ligating barcodes to libraries. Depending on the selected library preparation procedure, a number of quality controls and quantification measurements are performed. Reggie wizards are designed to provide configuration files for laboratory equipment and for parsing and reading generated results files throughout the process. Parsed data and results files are associated with library and bioplate items, representing used microtiter plates, respectively.

#### Create pooled libraries and Lab protocols for pooling

Wizards for creating pooled libraries are designed to take full advantage of having data on library concentration and size stored in BASE. After selecting a desired target pool concentration, the user can optimize pooling procedures by alternating between pooling strategies and using a fixed total volume or not. Depending on concentration of individual libraries, Reggie will automatically suggest omitting libraries with low yield from the pool and the user can manually overrule the suggestion and include or omit libraries freely as needed. Once pools are registered in Reggie, corresponding items are created in the database and lab protocols with full instructions for pooling are provided for printing.

#### Sequencing wizards

Wizards for sequencing are implemented to match the instrumentation used in the laboratory. For example, in earlier versions of Reggie, wizards for sequencing on the HiSeq Sequencing Systems (Illumina) were included, however, in the current implementation, wizards are restricted to support the NextSeq Sequencing Systems (Illumina). The restriction is simply an example of how development of Reggie follows operating procedures in the laboratory. Wizards for creating flow cells comprise an interface for registration of flow cells, flow cell type and flow cell identifier and for assigning a specific and available library pool to the flow cells. In earlier versions, where HiSeq flow cells are supported, multiple pools can be selected and assigned to individual lanes on the flow-cell. Lab-tracking sheets for clustering and sequencing, depending on Reggie version and flow cell type, are created and available for download by the user to assist with the practical work of clustering and configuring sequencing runs.

#### Register sequencing started and ended

Once a sequencing run is started, wizards are available to register the start and ultimately the end of the run. In these wizards, dialogs are available for specifying the location for data output on designated network file servers and for confirming sequencing parameters, such as number of cycles and type of sequencing done. Information on planned parameters, e.g., number of cycles, is already registered in the register flow cells wizard and defaults are configured to match operating procedures. However, used parameters are confirmed when the start of the sequencing run is registered. In addition, when the run has ended, Reggie verifies actual parameters by retrieving and parsing information from the files created by the sequencing machine. By designing Reggie wizards to continuously monitor the data output location, automatic detection and registration of a completed sequencing run can be triggered by specific output files from the sequencing machine. This in turn is utilized for automatic and prompt initiation of secondary analysis wizards that execute analysis pipelines for generated sequencing data.

#### Secondary analysis wizards

Secondary analysis includes several wizards for evaluating data files generated by the sequencing machine and for executing steps in the data analysis process. Individual steps can be initiated manually, however, wizards are implemented with optional auto-confirm functionality, permitting automatic registration and evaluation of completed analysis followed by initiation of subsequent steps. In this manner, automated analysis pipelines are made available for prompt completion of complex data processing and summarization of results. The implemented system is set up so that Reggie has access to a job scheduler to run jobs on external computation clusters, thereby reducing analysis time by supporting multiple parallel jobs to run simultaneously. To enable tracking of parameters used when running analysis tools, job parameters are stored in software and job items in the BASE database. Importantly, the auto-confirm functionality in Reggie includes rule-based verification of analysis, for example, an alignment is only auto-confirmed given that the aligned data fulfills required criteria such as a minimal number of aligned reads.

#### Confirm sequencing completed

After a sequencing run is registered as completed, Reggie will verify the presence of expected output files and that data has not become corrupt during writing or transfer to the file servers. If the data passes the integrity verification, automatic initiation of de-multiplexing and merge operations is done.

#### Start demux and merge

The implementation comprises integrated demultiplexing and merging, when libraries are multiplexed to multiple lanes or flow cells, followed by quality filtering. The process starts with demultiplexing using the external tool Picard ExtractIlluminaBarcodes and IlluminaBasecallsToFastq [13]. The wizard will automatically retrieve information on included libraries and used barcodes from BASE and supply it to Picard. After demultiplexing, data derived from individual libraries is analyzed in separate jobs, running Bowtie [14] for insert size estimation and Trimmomatic [15] for quality and adaptor filtering. After completion, individual derived bioassay items with associated fastq files containing read data are created. All relevant files are stored on file servers and URLs for these external objects are stored in BASE. Metainformation from the analysis, e.g., number of reads, number of adaptor reads, fragment size, and reads passing Trimmomatic is associated with the derived bioassays and stored in the database.

#### Start masking and alignment

Masking and alignment are executed as a single work-flow in Reggie and includes the creation of derived bioassay items with associated information and output files from alignment, e.g., BAM files with aligned reads. Masking is implemented by running Bowtie and removing reads that align to a masking reference sequence, e.g., including ribosome, PhiX, and standard repeats [7]. Reads that pass the masking continue through to alignment using Tophat [16] against a transcript and genome reference. After Tophat alignment is completed, Reggie runs Picard MarkDuplicates [13] to estimate the fraction of duplicate reads.

#### Start feature extraction

The Reggie wizard for feature extraction is implemented to run Cufflinks [17, 18] for estimating relative abundances of transcripts provided in a Gene transfer format (GTF) file. The Reggie work-flow runs Cufflinks on Tophat aligned reads, creates Raw bioassay items in BASE, and associates files from Cufflinks with these. In addition, the implementation parses Fragments Per Kilobase of transcript per Million mapped reads (FPKM) tracking files and imports expression data into BASE utilizing array designs as placeholders for features included in the transcript GTF file.

#### Report wizards

The implemented analysis pipeline includes wizards for summarizing data and creating reports that include RNAseq derived data. Since the extension package has access to all information related to the sequencing experiments, reports can include anything from patient information, biopsy properties, RNA quality scores, cellularity estimates from review of HE stained FFPE sections, as well as RNAseq gene expression estimates and biomarkers. Importantly, the system is implemented to create and store reports without sensitive personal patient information. A patient curator, logged in as a user with access to personal information, can access separate Reggie wizards to export clinical reports complete with added sensitive information.

These wizards also allow for downloading reports in password protected and strongly encrypted files for distribution to referring clinical departments.

#### Quality assurance, statistics and reporting wizards

A special set of wizards has been implemented to provide useful overall statistics and summaries of information stored in the database, e.g., number of enrolled patients stratified by referring hospital and year. An elaborate set of statistics can also be automatically compiled using wizards that plot sample processing performance over time. Such plots have proven useful to monitor yield, biopsy quantity, RNA quality, and more to highlight potential problems that might occur when otherwise unnoticeable changes are introduced, e.g., bad lot number for extraction columns or new routines at a referring pathology department.

In addition to the selection of Reggie wizards described herein, tools are included for administration and installing wizards and updates. Reggie has proven to be very flexible and as such has evolved significantly over time [19]. An important lead-time for a sequencing experiment is the time from acquiring the sample, to the time of completion of a set of analysis steps and ultimately a compiled report from sequencing data derived from the sample. Using BASE and extension package Reggie we have reduced this lead-time to less than 5 days, while still maintaining a sustainable, robust and less error prone workflow with extensive tracking of included laboratory and analysis steps. As of February 2016 more than 8000 enrolled breast cancer patients have been registered and are managed by the implemented system. Moreover, Reggie has managed sample processing for more than 6000 breast cancer biopsies, through partitioning, extraction, and library preparation, and the secondary analysis wizards have demultiplexed data from over 6000 sequenced libraries, executing the implemented analysis pipeline for masking, alignment, feature extraction, and creation of RNAseq reports. While the described implementation is designed to meet the need for a specific regional cancer genomics initiative, many of the implemented workflows in Reggie are designed for typical operating procedures used in many laboratories. The implementation can therefore be used as a template for creating a custom made implementation for any genomics project. The task in customization is to build a set of wizards similar to the ones we outlined above and that we collected into a single extension Reggie. For example, we have used the Reggie approach for implementing MeLuDI; an in-house project tailored to an initiative that concerns molecular diagnostics in melanoma and lung cancer (MeLuDI is available from the BASE plug-ins site [11]).

## Conclusions

Using free open-source software together with the latest generation BASE and customized extension package Reggie, we have successfully implemented a LIMS and data management system for a large-scale genomics project. With a high degree of automation and interactivity the system allows for resource efficient laboratory procedures and short lead-times which technically permit the use of RNAseq data analyses in clinical cancer management. The implementation described herein is designed to meet the need for a specific regional cancer genomics initiative but can be used as a template for creating a custom made implementation for any genomics project.

## Declarations

### List of abbreviations

BASE: BioArray Software Environment
SCAN-B: Sweden Cancerome Analysis Network - Breast
LIMS: Laboratory Information Management System
FFPE: Formalin-Fixed and Paraffin Embedded
HE: Hematoxylin and Eosin
GTF: Gene Transfer Format
FPKM: Fragments Per Kilobase of transcript per Million mapped reads
XML: Extensible Markup Language

## Availability of software and supporting information

BASE is a web application residing in a Tomcat container and Reggie is an add-on to BASE. Both BASE and Reggie are written in the Java programming language and requires Java 8 or later and Apache Tomcat 8. BASE is operating system agnostic and runs on any operating system that supports Java 8 and Apache Tomcat 8.

BASE and extension package Reggie are available as open-source software. BASE is released under the GNU General Public License and is available for download from the BASE project site (http://base.thep.lu.se). Reggie is released under GNU Affero General Public License and is available for download from the BASE plug-ins site (http://baseplugins.thep.lu.se).

## Authors' contributions

JH, NN, and JVC designed BASE and the Reggie extension. JH and JVC integrated BASE and Reggie within the laboratory. JH coordinated the BASE project. NN was the lead software developer and implemented BASE and Reggie. OM implemented LabEnv and assisted in Reggie implementation. JVC managed project development and wrote the manuscript. All authors read and approved the final manuscript.

## Acknowledgements

We thank the personnel of the SCAN-B laboratory for comments and feedback throughout the process of implementing the system and integrating BASE and Reggie within the laboratory.

